# Hingepoint emergence in mammalian spinal neurulation

**DOI:** 10.1101/2021.10.22.465433

**Authors:** Veerle de Goederen, Roman Vetter, Katie McDole, Dagmar Iber

**Affiliations:** Department of Biosystems Science and Engineering (D-BSSE), ETH Zürich, 4058 Basel, Switzerland; Graduate School of Life Sciences, Utrecht University, 3584CG Utrecht, Netherlands; Department Health Sciences and Technology (D-HEST), ETH Zürich, 8603 Schwerzenbach, Switzerland; Swiss Institute of Bioinformatics (SIB), 4058 Basel, Switzerland; MRC Laboratory of Molecular Biology, Cambridge Biomedical Campus, Cambridge CB2 0QH, United Kingdom

## Abstract

Neurulation is the process in early vertebrate embryonic development during which the neural plate folds to form the neural tube. Spinal neural tube folding in the posterior neuropore changes over time, first showing a median hingepoint, then both the median hingepoint and dorsolateral hingepoints, followed by dorsolateral hingepoints only. The biomechanical mechanism of hingepoint formation in the mammalian neural tube is poorly understood. Here, we employ a mechanical finite element model to study neural tube formation. The computational model mimics the mammalian neural tube using microscopy data from mouse and human embryos. While intrinsic curvature at the neural plate midline has been hypothesized to drive neural tube folding, intrinsic curvature was not sufficient for tube closure in our simulations. We achieved neural tube closure with an alternative model combining mesoderm expansion, non-neural ectoderm expansion and neural plate adhesion to the notochord. Dorsolateral hingepoints emerged in simulations with low mesoderm expansion and zippering. We propose that zippering provides the biomechanical force for dorsolateral hingepoint formation in settings where the neural plate lateral sides extend above the mesoderm. Together, these results provide a new perspective on the biomechanical and molecular mechanism of mammalian spinal neurulation.

## 1 Introduction

Neurulation is the process in early chordate embryogenesis during which the neural tube (NT) forms. The neural tube is a dorsal structure extending along the rostrocaudal axis which ultimately develops into the brain and spinal cord. Failure of neurulation leads to neural tube defects, a group of severe neurodevelopmental disorders that affect approximately 1 in 1,000 births [1]. Neurulation occurs through two distinct mechanisms termed primary and secondary neurulation. In amphibians, birds, and mammals, primary neurulation (reviewed in [2]) occurs along most of the rostrocaudal axis, whereas secondary neurulation (involving epithelialization of condensed tail-bud cells) occurs caudally in the lower sacral and coccygeal regions only. At the start of primary neurulation, dorsal midline ectoderm cells differentiate into neuroepithelial cells to form a columnar pseudostratified epithelium called the neural plate (NP) (reviewed in [3]). The NP lateral sides elevate and meet dorsally at the embryo midline, where they fuse to form the NT. The folding NP is bordered by non-neural ectoderm (NNE) cells, which adhere to the neural plate borders (NPBs) basal side and migrate medially to cover the NT at the dorsal midline [2, 4].

While amphibians show neural tube closure at all axial levels simultaneously, mammals and birds initiate closure at a few sites, with the exact closure locations differing between species [2]. In mice, the neural tube initially closes on embryonic day (E) 8, somite stage (SS) 7, at the hindbrain/cervical boundary between somite 3 and 4 (closure 1, C1) [5, 6]. Starting from the initial closure site, neural tube closure progresses bidirectionally through “zippering” (Fig. 1A). During zippering, the NNE pulls the NPBs towards the embryo midline by forming a transitory semi-rosette structure at the dorsal closure site [7]. The NP region caudal of the “zipper” is called the posterior neuropore (PNP) [2] (Fig. 1A, close-up box). Simultaneous with zippering progression, the PNP expands in caudal direction until complete closure is achieved at E10, SS30 [5].

**Figure 1:**
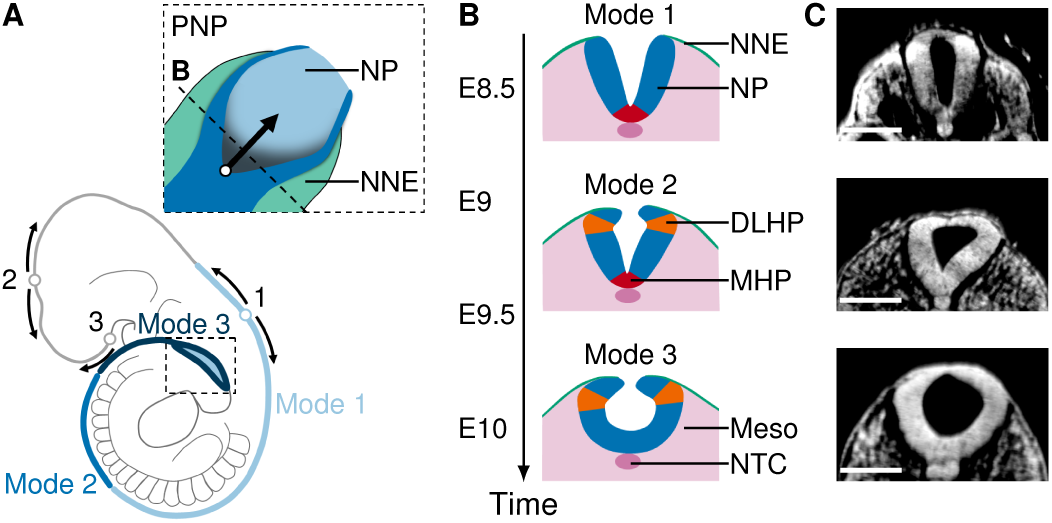
Neural plate morphology in the posterior neuropore at successive stages of mouse neurulation. (A) Schematic mouse embryo at E10. Dotted box, close-up of PNP (top view); empty circles, closure sites (closure 1-3); arrows, zippering direction; blue shades, position of folding modes 1–3. (B) Schematic transverse sections through the PNP at spinal neurulation modes 1 (E8.5–E9), 2 (E9–E9.75) and 3 (E9.75–E10.5). Green, NNE; blue, NP; red, MHP; orange, DLHPs; pink, mesoderm (Meso) and notochord (NTC). (C) Transverse sections through the PNP at E8.5 (Mode 1), E9.5 (Mode 2) and E10 (Mode 3). Scale bars: 100 µm. Images were derived from [8] under a Creative Commons Attribution license.

When looking at transverse cross-sections of the PNP, the folding NP shows strong curvature in two specific regions, commonly referred to as “hingepoints”. Bending occurs at the ventral midline, resulting in the median hingepoint (MHP), and at lateral sides near the position where the NP touches the NNE, resulting in the dorsolateral hingepoints (DLHPs) [4, 9]. Interestingly, the spinal neural tube appears to fold through different mechanisms along the rostrocaudal axis. At E8.5, when the upper spine forms, the folding neural tube shows a MHP but no DLHPs (Mode 1). At E9.5, during mid-spine formation, the neural tube shows both a MHP and DLHPs (Mode 2). When the posterior neural tube closes at E10, only DLHPs are present (Mode 3) (Fig. 1B,C) [9]. Mouse mutants with impaired MHP formation show increased DLHPs [10], and mouse *Noggin*^-/-^ mutants, which fail to form DLHPs, show NT closure failure in the most caudal regions [11, 12]. While hingepoints appear to be required for tube closure, the biomechanical and molecular mechanism of MHP and DLHPs formation is poorly understood.

Hingepoints are commonly thought to form through an intrinsic curvature mechanism involving apical constriction and basal widening of neuroepithelial cells. Avian and mammalian neuroepithelial cells are tightly packed in a columnar pseudostratified epithelium, such that the cells are widened at the apicobasal position of the nucleus. Neuroepithelial nuclei move along the apicobasal axis throughout the cell cycle in a process termed interkinetic nuclear migration (IKNM, reviewed in [13]). While most neuroepithelial cells are distributed randomly throughout the cell cycle, the MHP contains an increased proportion of wedge-shaped S-phase cells with basally located nuclei. This observation led to the view that IKNM is a driving force in avian [14] and mammalian [15, 16] MHP formation. However, while DLHPs show an increased proportion of cells with basally located nuclei in the chick [14], no correlation has been found in mice [16]. Furthermore, disruption of actin microfilaments does not prevent hingepoint formation in the mouse spinal NP, contradicting the idea that hingepoints form through cell-intrinsic bending [17].

Aside from intrinsic forces of NP bending, extrinsic forces from the notochord, mesoderm and NNE have been implicated in NTF. The NP adheres to the notochord [15], and the notochord is essential for MHP formation. Notochordless embryos lack strong midline curvature and do not develop a floor plate [10, 18, 19]. In addition to the notochord, mesoderm expansion has been proposed to drive neural tube folding in the rostral region [20, 21]. Neural tube closure at the lower spinal cord region requires presence of NNE, but can still form after mesoderm removal [22, 23].

Computational modeling allows for an isolated study of biomechanical NTF driving forces in a defined and quantifiable setting. Previous modeling studies have focused on amphibian neurulation [24–27]. Due to differences in closure progression and NP morphology, a new model is needed which is specific for mammals. Here, we employ a 2D finite element model to study the biomechanics of NTF. We find that mesoderm and NNE expansion combined with NP-notochord adhesion are sufficient to simulate MHP formation and Mode 1 closure. Furthermore, we find that DLHPs emerge in simulations with low mesoderm expansion and zippering.

## 2 Results

### Intrinsic neural plate curvature and midline tapering can promote tube folding

To investigate the mechanical mechanism of neural tube folding, we first examined the minimal requirements to form a tube from a flat tissue. In order for an epithelial sheet covering an embryo to fold into a tube structure, it is necessary that the epithelial tissue expands relative to the underlying tissue volume (differential growth). In concordance, we modeled an expanding epithelial 2D section with spatially fixed boundaries, dubbed “Model I”.

The initial Model I configuration is a horizontal tissue of width *w*_NP_ = 500 µm and apicobasal height *h*_NP_ = 45 µm (Fig. 2A, simulation time *t* = 0). The ectoderm expands in width uniformly over space and time, whereas the apicobasal height remains constant. The simulation ends at twice the initial tissue length (1000 µm, simulation time *t* = 1). In order to mimic differential growth of the ectoderm with respect to underlying tissues, the net area under the ectoderm horizontal midline is kept at zero using a Lagrange multiplier. As deforming embryonic tissues show viscoelastic properties [28], we modeled the epithelium as a linearly viscoelastic continuum. Furthermore, as biological cells show stress relaxation in response to external forces through cytoskeletal remodeling [29], we included plastic stress relaxation in the model (see Materials and Methods). For an introduction to tissue mechanics, we refer to [30].

**Figure 2:**
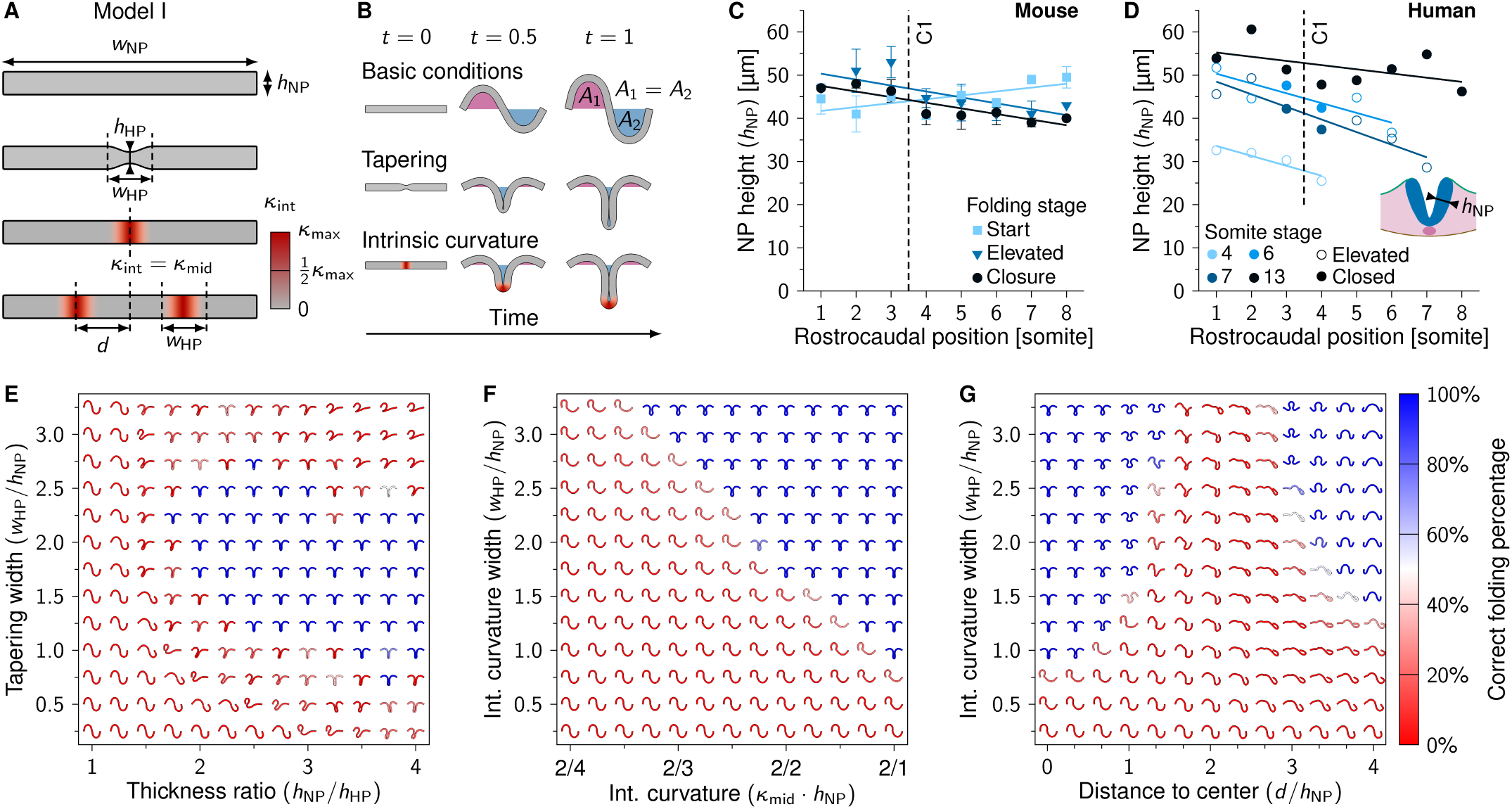
Intrinsic neural plate curvature and midline tapering can promote tube folding. (A) Schematic representation of Model I. *w*_NP_, neural plate width; *h*_NP_, neural plate apicobasal height; *h*_HP_, median hingepoint (midline) height; *w*_HP_, hingepoint domain width; *d*, domain distance to center; *κ*_int_, intrinsic tissue curvature; *κ*_mid_, intrinsic tissue curvature at the hingepoint midline; *κ*_max_, theoretical maximum tissue curvature. (B) Model I simulations rendered between *t* = 0 (*w*_NP_ = 500 µm) and *t* = 1 (*w*_NP_ = 1000 µm). The area underneath the tissue horizontal midline is fixed at 0. (C) Measured neural plate apicobasal height (*h*_NP_) during neural tube folding progression along the rostrocaudal axis in developing mouse embryos. Error bars indicate SEM, lines represent linear least-squares fit, C1 indicates closure 1 rostrocaudal position. *N* = 1–3 (see Table S1). (D) Measured neural plate apicobasal height (*h*_NP_) along the rostrocaudal axis in human embryos (SS4, SS6, SS7 and SS13). Lines represent linear least-squares fit. Open circles, NP is in the process of folding; closed circles, NP is closed. (E-G) Neural plate folding simulations (*t* = 1) with midline tapering thickness ratio versus tapering domain width (E), intrinsic midline curvature strength versus intrinsic curvature domain width (F) and intrinsic dorsolateral curvature (*κ*_mid_ = *κ*_max_*/*2) distance to center versus intrinsic curvature domain width (G). In (E), simulation renderings were flipped along the *y*-axis to point downwards. Color scale represents the percentage of intermediate simulation steps in which the neural ectoderm is sufficiently symmetrical along the *x*-axis and the neuroepithelium midline is buckled, i.e., the sum of the x-coordinates of the elements on the left and right half of the neural ectoderm differ by less than 10%, and the middle element has |*y*| > *h*_NP_.

Epithelial width expansion resulted in tissue outgrowth, but did not lead to tube formation (Fig. 2B top row, Movie S1). We subsequently tested the effect of midline tapering on neural plate folding. Neuroepithelial cells at the NP midline have a lower apicobasal height than lateral neuroepithelial cells, due to SHH signaling from the notochord [18, 19, 31]. Bending energy scales with the tissue thickness cubed [32], such that tissues subject to transverse forces tend to bend at the thinnest sites. Thus, midline tapering may facilitate MHP formation and tube folding. We implemented midline tapering in Model I using a thickness ratio given by the lateral neural plate height (*h*_NP_) divided by the midline neural plate height (*h*_HP_) (Fig. 2A). To determine lateral neural plate height, we analyzed *in vitro* live imaging data of three developing mouse embryos expressing a cell membrane and nuclear marker (E6.5–E8.5, Fig. 2C). We analyzed one of our previously published mouse embryo data sets [33] and combined this with two new ones. To compare mouse with human NTF, mouse live imaging data was complemented with stained histological sections of four human embryos between SS4–SS13 (Fig. 2D). Human embryo sections were obtained from the Virtual Human Embryo Project [34] and the 3D Atlas of Human Embryology [35]. Mouse and human NP height both showed a combined mean of 44 µm measured across samples and rostrocaudal positions (44*±*5 µm and 44*±*8 µm (SD), respectively), which we rounded off to *h*_NP_ = 45 µm for the simulations. We modeled the neural epithelium height transitions from *h*_HP_ to *h*_NP_ using a smooth function covering the median hingepoint domain width (*w*_HP_) (Fig. 2A). We ran Model I simulation for varying thickness ratios and tapering widths. Tubes with left-right symmetry formed at thickness ratios of 1.75 and higher and at tapering widths between 0.75 and 2.75 times the NP height (Fig. 2E, Movie S2). As midline thinning was symmetrical along the apicobasal axis, tubes folded in both outward and inward direction. Thus, while our simulation results suggest midline tapering could promote tube folding, an additional process is needed to ensure the tube folds inward.

As a possible mechanism to ensure inward folding, we next considered intrinsic midline curvature. NTF is associated with increased presence of wedge-shaped cells at the MHP [15, 16]. The formation of wedge-shaped cells with narrow apical and wide basal sides is considered a driving force of NTF [2, 15, 16]. We ran Model I simulations for varying levels of intrinsic midline inward curvature and MHP width values (Fig. 2A,B,F, Movie S3). Intrinsic curvature strength is given by *κ*_int_ = 1*/r*_c_, with *r*_c_ the radius of curvature. As the theoretical minimum radius of curvature corresponds to the tissue half-height (*h*_NP_*/*2), the theoretical maximum intrinsic curvature *κ*_max_ = 2*/h*_NP_. We modeled intrinsically curved regions using a spatially smooth and temporally instantaneous function ranging from *κ*_int_ = *κ*_mid_ in the middle of the intrinsically curved domain to *κ*_int_ = 0 on the lateral domain borders. Note that simulations with intrinsic curvature start out in a straight configuration; the intrinsically curved regions curve up during early simulation time steps (Fig. 2A,B). Intrinsic midline curvature combined with NP width expansion lead to tube formation for intrinsic curvature domain widths *w*_HP_ = *h*_NP_ and higher, and intrinsic curvatures above *κ*_mid_ = *κ*_max_*/*3.5 (Fig. 2F). Wide intrinsically curved domains were favorable for folding and required weaker intrinsic curvature to form a neural tube. In contrast to midline tapering, intrinsic midline curvature was biased in the ventral direction and thus consistently lead to inward folding.

As intrinsic midline curvature caused a tube shape with MHP resembling Mode 1 folding (E8.5), we wondered if likewise we could simulate a tube with DLHPs resembling Mode 3 (E10) using intrinsic dorsolateral curvature. To implement this, we introduced the parameter *d* to indicate the distance between the NP midline and the middle of the DLHP intrinsic curvature domains (Fig. 2B, bottom row). Intrinsic dorsolateral curvature was fixed at *κ*_mid_ = *κ*_max_*/*2 and distance *d* was varied from 0 (hingepoints overlap at the midline) to 180 µm (hingepoints positioned close to NPBs). For low *d*, the intrinsically curved regions overlapped to form a median hingepoint. However, when the intrinsically curved regions were shifted laterally to form separate DLHPs (at *d* = 75–120 µm), the epithelium folded into an irregular shape not resembling a tube (Fig. 2G, Movie S4). These results suggest that epithelium width expansion combined with intrinsic neural plate curvature may be sufficient for tube formation when the neural plate curves in the midline region, but not when the neural plate curves in dorsolateral regions.

When the inward DLHP intrinsic curvature domains approached the NPBs (*d* > 120 µm), a tube shape did form, albeit consistently in the outward direction (Fig. 2G). Thus, curvature at the NPBs in the outward direction could contribute to tube folding. Such curvature has been commonly observed in birds and mammals, and has previously been described as “epithelial ridging and kinking” at the NPBs as an event preceding adhesion between the NNE and NP basal sides [36].

### Non-neural ectoderm width increase in neural tube folding

After studying intrinsic neural plate curvature and midline tapering in a minimal model, we adapted the model to better represent mammalian NTF. NP width decreased throughout folding by an average of 30% in live-imaged mouse embryos (360±73 µm at folding start versus 251±42 µm (SD) at time of closure) whereas no clear trend was observed in human embryos (Fig. 3A,B). The NP is bordered by NNE on the lateral sides, which increased in width throughout folding (48% increase from 175±46 µm at folding start versus 259±29 µm (SD) at time of closure in mice) (Fig. 3C,D). The NNE apicobasal height was measured at 7.2±1.2 µm (SD) in human embryos, which is approximately 6 times shorter compared to the NP (Fig. 3E). According to linear elastic beam theory [32], this makes NNE 6^3^≈216 times less rigid in bending than the NP. After incorporating this information into Model I, midline tapering (with *h*_NP_*/h*_MHP_ = 2, *w*_HP_ = 2*h*_NP_) and midline intrinsic curvature (*κ*_mid_ = *κ*_max_, *w*_HP_ = 2*h*_NP_) were no longer sufficient to create a tube shape (Fig. 3F). When combining midline tapering and maximum midline intrinsic curvature in the original Model I, a tube formed (Fig. 3F, top row, Movie S5). However, the folding neural plate did not close after incorporating the NNE thickness profile and differential width expansion in the model, with a fixed *w*_NP_ and only *w*_NNE_ expanding over time (Fig. 3F, second and third row, Movies S6 and S7). Even when adding maximum outward intrinsic curvature at the NPBs to promote NP elevation (*κ*_mid_ =−*κ*_max_, *w*_HP_ = 2*h*_NP_, *d* = *w*_NP_*/*2), the NP did not close (Fig. 3E, bottom row, Movie S8). While in theory we could force the NP to form a tube by increasing the intrinsic curvature domain width, this is not consistent with observations from birds and mammals, which during Mode 1 folding only show sharp bending at the NP midline [14–16]. Together, these results suggest that intrinsic NP midline curvature and midline tapering are not sufficient for NT closure in mammals and birds. Thus, an additional or different mechanism is needed.

**Figure 3:**
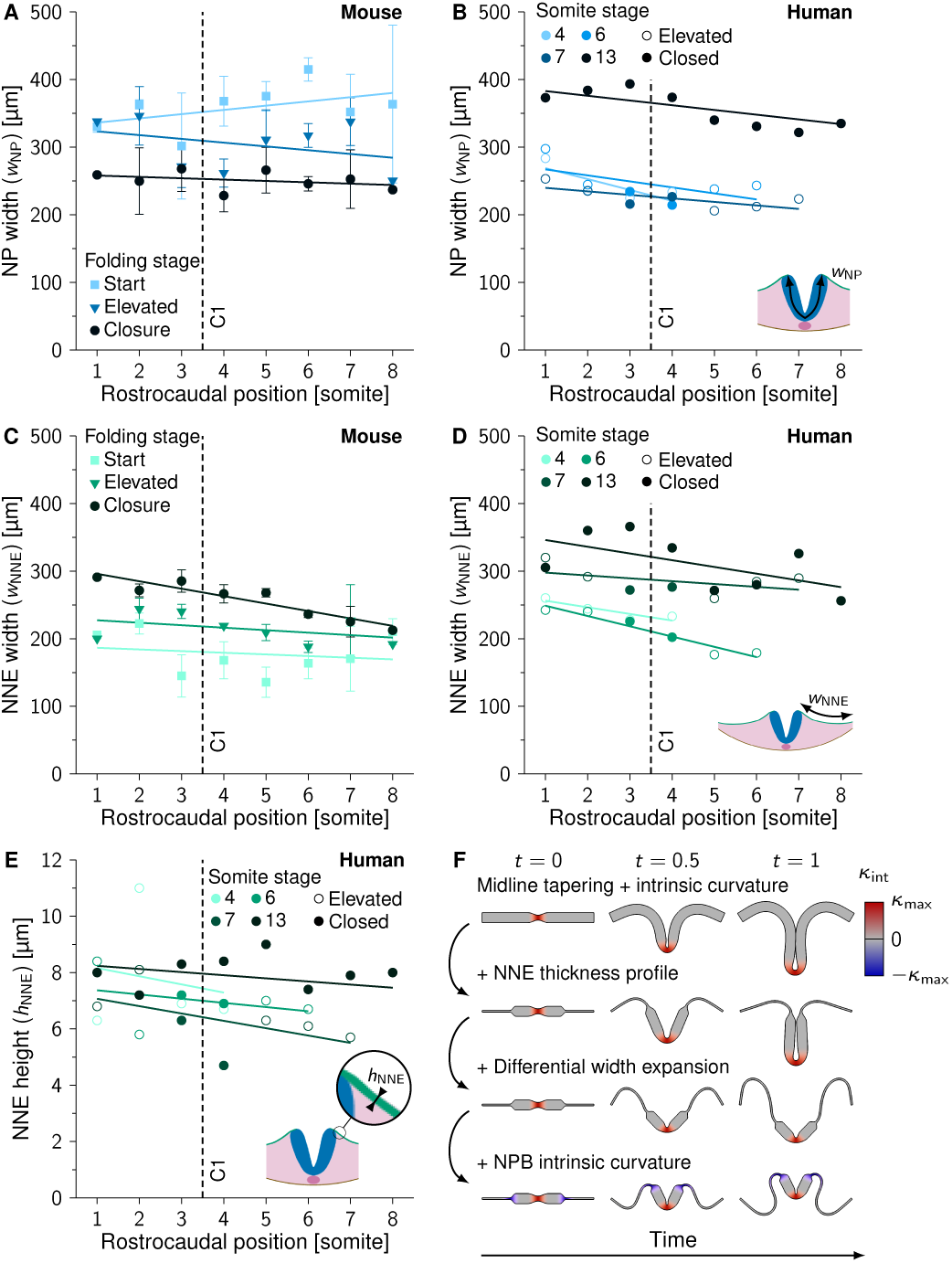
Non-neural ectoderm width expansion in neural tube folding. (A) Measured NP width (*w*_NP_) during neural tube folding progression along the rostrocaudal axis in developing mouse embryos. *N* = 1–3 (see Table S1). (B) Measured NP width (*w*_NP_) along the rostrocaudal axis in human embryos (SS4, SS6, SS7 and SS13). (C) Measured NNE width (*w*_NNE_) during neural tube folding progression along the rostrocaudal axis in developing mouse embryos. *N* = 1–3 (see Table S1). (D) Measured NNE width (*w*_NNE_) along the rostrocaudal axis in human embryos (SS4, SS6, SS7 and SS13). (E) Measured NNE height (*h*_NNE_) along the rostrocaudal axis in human embryos (SS4, SS6, SS7 and SS13). Lines in (A–E) represent linear least-squares fits, C1 in (A–E) indicates closure 1 rostrocaudal position, error bars in (A, C) indicate SEM. (B, D, E) open circles, NP is in the process of folding; closed circles, NP is closed. (F) Model I simulations rendered between *t* = 0 and *t* = 1. Top row, midline thinning + midline intrinsic curvature; second row, previous simulation + NNE thickness profile; third row, previous simulation + differential width expansion; bottom row, previous simulation + NPB intrinsic curvature. Red, inward (midline) intrinsic curvature; blue, outward (NPB) intrinsic curvature.

### Extrinsic forces are sufficient for neural tube closure

As intrinsic neural plate curvature appeared insufficient for mouse neural tube closure, we next considered extrinsic forces. Mesoderm tissue underlying the NP increased by 58% in area during mouse NTF (1.9*×*10^4^*±*0.4*×*10^4^ µm^2^ at folding start versus 3.0*×*10^4^*±*0.6*×*10^4^ µm^2^ (SD) at time of closure) (Fig. 4A). While no such trend was found for the human embryo sections, this can likely be attributed to size variations between individual embryos (Fig. 4B). Mesoderm expansion has been proposed to promote NP elevation [20, 21]. Importantly, the neural plate midline adheres strongly to the underlying notochord [15], thus possibly anchoring the NP midline while the lateral sides elevate. In addition to the notochord and mesoderm, NNE has been implicated in NTF [22, 23]. The NNE expanded in width during mouse NTF (Fig. 3C). While the NNE appeared too thin to push the neural folds towards the midline in Model I (Fig. 3F), NNE could nevertheless promote NTF in a biomechanical model by constraining the direction of mesoderm expansion.

**Figure 4:**
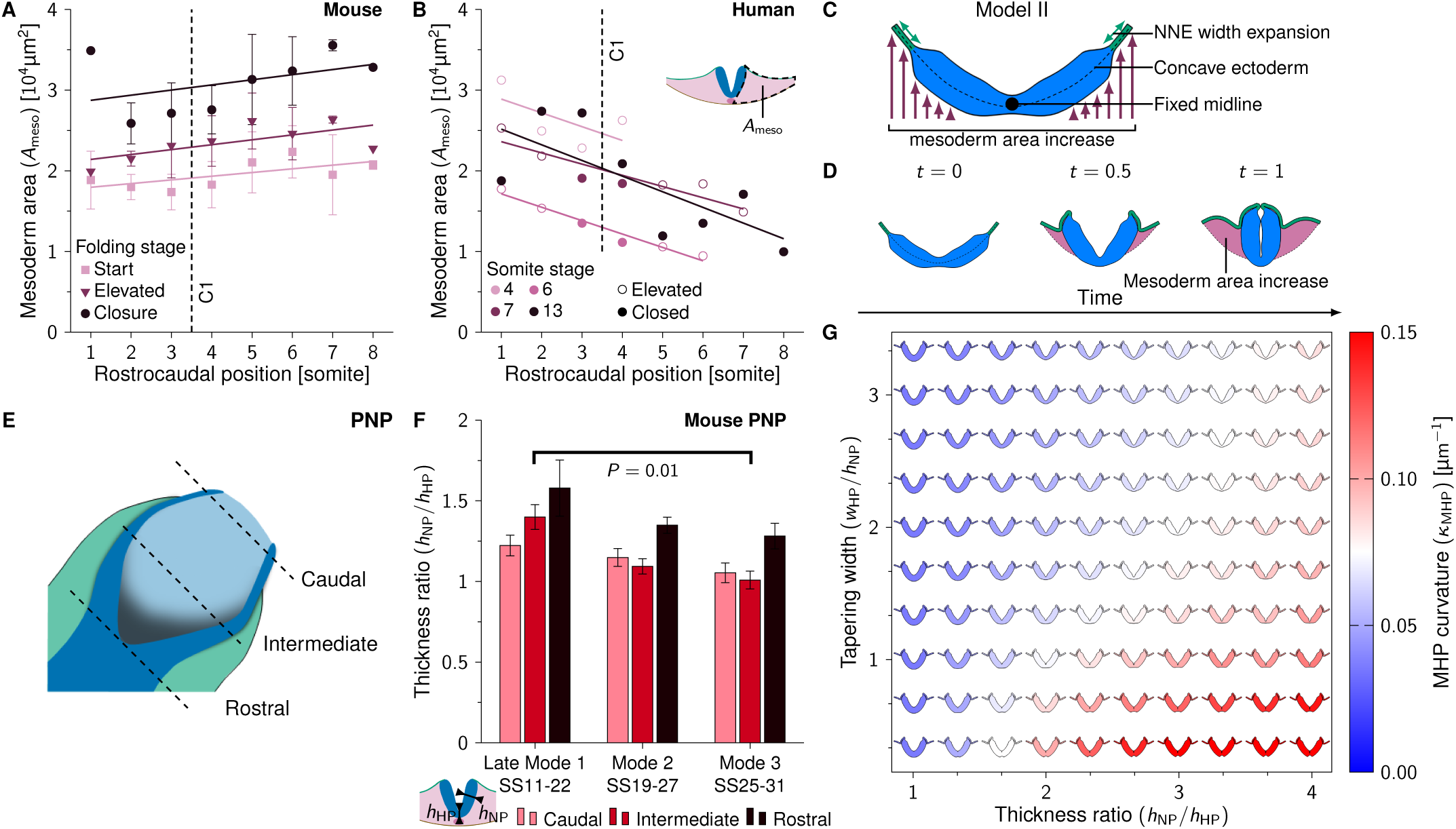
Mesoderm area increase and midline tapering promote MHP formation. (A) Measured mesoderm area (*A*_meso_) during neural tube folding progression along the rostrocaudal axis in developing mouse embryos. *N* = 1–3 (see Table S1). Error bars indicate SEM, lines represent linear least-squares fits. (B) Measured mesoderm area (*A*_meso_) along the rostrocaudal axis in human embryos (SS4, SS6, SS7 and SS13). Open circles, NP is in the process of folding; closed circles, NP is closed. Lines represent linear least-squares fits. (C) Schematic representation of Model II. The NP midline position is held fixed. The NNE width expands over time and the area ventral of the ectoderm increases to simulate mesoderm expansion. The initial ectoderm configuration is concave. (D) Model II simulation rendered between *t* = 0 and *t* = 1. The pink area indicates mesoderm area increase (1.7*×*10^4^ µm^2^ at *t* = 1). (E) Schematic drawing of the PNP. Caudal, NP is flat; Intermediate, NP is elevated; Rostral, NP closes. (F) Measured thickness ratio (*h*_NP_*/h*_HP_) during neural tube folding progression at Mode 1–3 in the developing mouse PNP. Late Mode 1, *N* = 5; Mode 2, *N* = 12; Mode 3, *N* = 7. Mann–Whitney *U* test, *P* = 0.01. (G) Model II simulations (*t* = 0.5) with midline tapering ratio versus median hingepoint domain width. Color scale indicates the highest curvature measured in the MHP region (*κ*_MHP_).

We combined mesoderm area increase, NNE width increase and NP-notochord adhesion into a new model (Model II, Fig. 4C). Unlike Model I, Model II does not keep the area underneath the ectoderm fixed but rather lets it increase linearly over time to simulate mesoderm area increase. The NNE simultaneously expands in width to facilitate the increasing mesoderm area. The NP remains constant in height and width during folding progression. To simulate notochord adhesion, the NP midpoint position is held fixed. To mimic the E8.5 embryo shape during early NT closure (Movie S9), we started the simulation from a concave ectoderm shape. Furthermore, to create more realistic ectoderm behavior we added adhesion between the NNE and NP basal sides upon reaching a proximity of 3 µm. In the embryo, neuroepithelial thickening during NP formation promotes NP-NNE adhesion [37]. To facilitate initial NP-NNE adhesion in the model, we included intrinsic outward curvature at the NPBs (*κ*_mid_ = *κ*_max_, *w* = 3*h*_NNE_). When combining mesoderm area increase and NNE width increase in Model II, the NP lateral sides elevated and moved towards the embryo midline to form a tube (Fig. 4D, Movie S10). Note that Model II is based on external forces and thus does not rely on intrinsic NP midline curvature. This result suggests that mesoderm and NNE expansion combined with notochord adhesion are sufficient for neural tube closure in the concave-shaped E8.5 embryo.

### Midline tapering enhances MHP curvature

In Model II (Fig. 4D), the MHP emerges through mesoderm expansion and NP midline adhesion to the notochord. During folding, the PNP shows a gradual decrease in midline curvature over time until there is no distinct MHP in the mouse PNP at E10 (Fig. 1B). To understand these observed differences along the rostrocaudal axis, we studied which factors determine MHP curvature. As elastic structures tend to deform at their thinnest point, where least force is required [32], we considered midline tapering. To see how midline tapering changes over developmental time between Mode 1 and Mode 3 folding, we analyzed high-resolution plastic sections and HREM data of the mouse PNP between Modes 1–3 (SS11–SS31). We refer to the PNP Mode 1 data (SS11–SS22) as “late Mode 1” to distinguish it from the early Mode 1 folding in our mouse live imaging data (SS8), as we observed relevant morphological differences between these time points (to be discussed in the next section). Mouse embryo images were obtained from [38] and [8]. We took transverse sections of the PNP at the caudal (NP is flat), intermediate (NP is elevated) and rostral (NP closes) sites (Fig. 4E). While Mode 1 folding showed midline tapering in the elevated PNP (thickness ratio 1.4*±*0.1 (SD)), no midline tapering was present in the elevated PNP at Mode 3 (thickness ratio 1.0 *±* 0.1 (SD)). No significant changes in midline tapering were found at the rostral and caudal PNP. Next, we ran Model II simulations for varying midline thickness ratios and midline tapering widths. We measured the curvature in the MHP region at simulation halftime *t* = 0.5. Higher thickness ratios promoted midline curvature (Fig. 4G). Interestingly, simulations without midline thinning showed a rounded NP midline that looked similar to the *Shh* mouse mutant, which lacks floor plate development (Movie S11) [10, 31]. Furthermore, midline curvature decreased with increasing tapering widths. Together, these results indicate that high NP midline thickness ratios and low tapering widths can enhance mesoderm-driven MHP curvature.

### DLHPs emerge through zippering at sites with low mesoderm expansion

After studying MHP formation, we sought to identify the biomechanical driving force of DLHPs. DLHPs form in the PNP at the 19–30 somite stage [10]. As the PNP caudal NP is flat rather than concave, we adapted Model II to start out in a horizontal configuration. To obtain a simple relationship between mesoderm area increase and NNE width increase, we approximated the growing embryo as a semiellipse, introducing the linear mesoderm area increase factor *α. α* = 1 corresponds to the area of a semiellipse with the lateral simulation boundary separation as the lateral diameter and *w*_NP_ as the dorsoventral diameter. The mesoderm area was increased linearly over time and the NNE width was increased linearly over space and time to match the area and corresponding circumference of the semicircle at the end of the simulation (*t* = 1). Model II simulations with *α* = 1 showed NP elevation, but mesoderm area expansion was not sufficient to reach NT closure (Fig. 5A, top row, Movie S12). When increasing mesoderm area expansion (*α* = 1.75, NNE width increase unchanged), the folding NP was forced to close. However, closure was initiated at the mid-lateral sides, rather than at the NPBs, and the resulting mesoderm area was significantly larger compared to the mesoderm area in the PNP at NT closure (Fig. 5A, middle row, Movie S13). As PNP closure depends on zippering [39], zippering could function to ensure closure at sites with insufficient mesoderm area expansion. To incorporate zippering in the simulations, we fixed the NP borders on a linear trajectory along the *x*-axis, such that the NP borders reach the embryo midline at *t* = 1. Simulating zippering (*α* = 1) lead to tube formation with a slit-shaped lumen characteristic of Mode 1 closure (Fig. 5A, bottom row, Movie S14).

**Figure 5:**
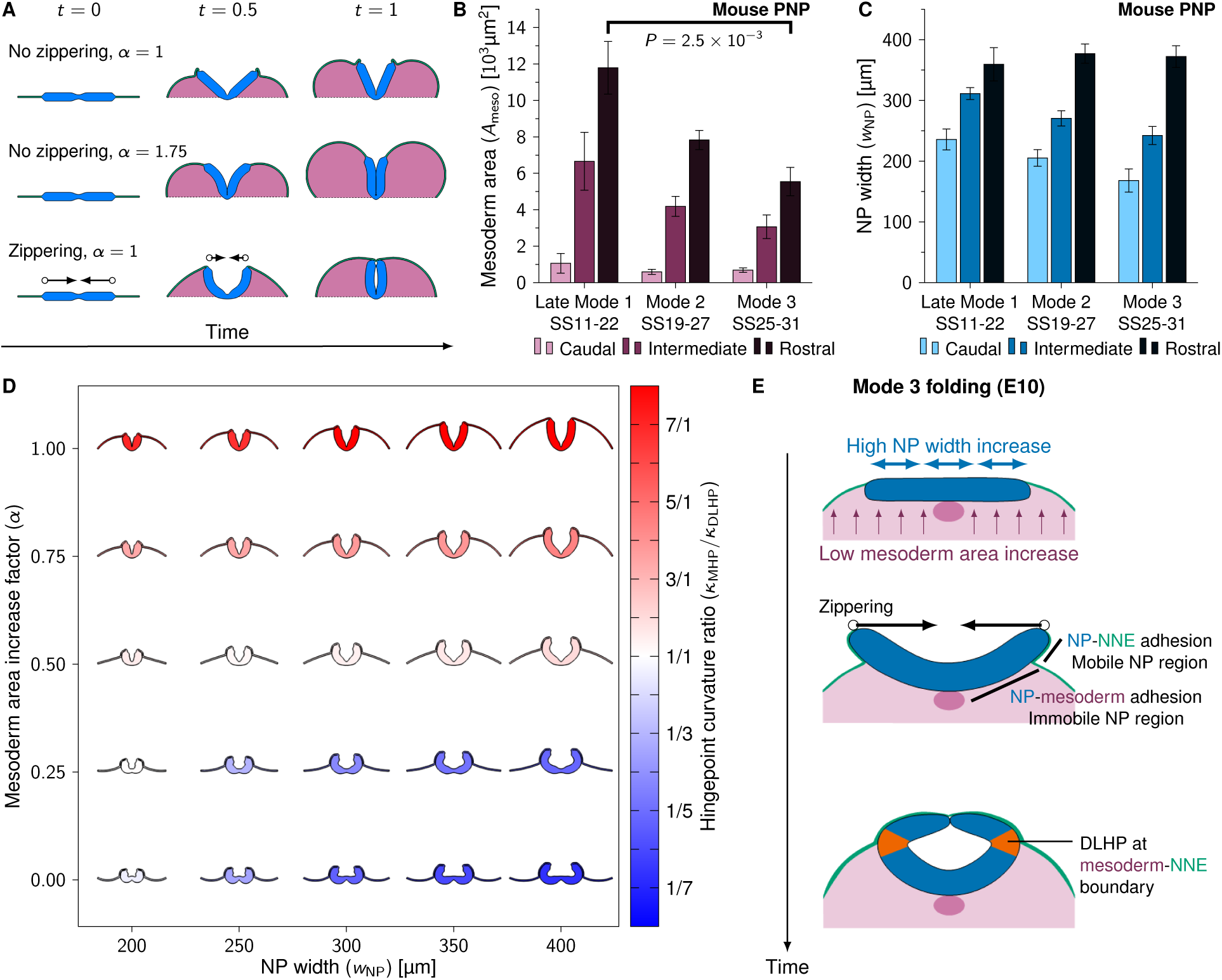
Zippering drives DLHP formation at sites with low mesoderm area increase. (A) Model II simulations with flat initial configurations rendered between *t* = 0 and *t* = 1. Arrows indicate zippering direction, pink area indicates mesoderm area increase. (B, C) Measured mesoderm area (*A*_meso_, B) and NP width (*w*_NP_, C) during neural tube folding progression at Modes 1–3 in the developing mouse PNP. Late Mode 1, *N* = 5; Mode 2, *N* = 12; Mode 3, *N* = 7. Error bars indicate SEM. (B) Mann–Whitney *U* test, *P* = 2.5*×*10^−3^. (D) Neural plate folding simulations (Model II + zippering) rendered at *t* = 0.7 with NP width (*w*_NP_) versus mesoderm area increase factor (*α*). Color indicates the ratio of MHP curvature versus DLHP curvature. Red, MHP dominates (Mode 1); white, both MHP and DLHPs (Mode 2); blue, DLHPs dominate (Mode 3). DLHP curvature was the highest curvature measured between the MHP and NPB regions at one of the NP lateral sides. (E) Proposed biomechanical mechanism of DLHP formation. The E10 PNP shows high NP width increase and low mesoderm area increase (top panel), which causes the NP lateral sides to elevate above the mesoderm and form basal contact with the NNE (middle panel). The lateral NP regions are therefore mobile and get pulled inwards through zippering, resulting in DLHPs at the mesoderm-NNE boundary (bottom panel).

To identify a potential biomechanical mechanism of DLHP formation, we analyzed high resolution plastic sections and HREM data of the mouse PNP between late Mode 1 and Mode 3 (SS11–SS31). The progression from late Mode 1 (no DLHPs) to Modes 2/3 (DLHPs) showed a strong reduction in mesoderm area lateral to the NP at the rostral PNP, with Mode 3 showing over 45% reduction in mesoderm area compared to late Mode 1 (Mann–Whitney *U* test, *P* = 2.5*×*10^−3^) (Fig. 5B). For NP width we found no significant differences between late Mode 1 and Modes 2/3, with and average width of 360 ± 6 µm (SD) at the closure site in the rostral PNP (Fig. 5C). Note however that the late Mode 1 NP width dynamics differ significantly from early Mode 1 folding (somite position 1–8) as observed in our live-imaging data (Fig. 3A). While the NP decreased by 30% in width during early Mode 1 folding to reach 251*±*42 µm (SD) at time of closure (Fig. 3A), late Mode 1 folding showed a 53% increase in NP width from 236 *±* 38 to 360 *±* 6 µm (SD) between caudal and rostral PNP, which can be considered a proxy for closure progression over time.

To test whether mesoderm expansion and NP width influence DLHP formation, we ran Model II simulations with zippering for varying levels of mesoderm area expansion (*α* between 0 and 1) and varying NP widths (*w*_NP_ between 200 and 400 µm). Strikingly, DLHPs emerged in simulations with decreased mesoderm expansion (Fig. 5D, Movies S15 and S16). Moreover, DLHPs became more visible at higher NP widths. Together, these results indicate that DLHPs could emerge in the late-stage PNP as a result of zippering in areas with low mesoderm expansion and high NP width.

## 3 Discussion

We combined mathematical modeling and quantitative image analysis to explore the role of mechanical forces in mammalian spinal neural tube folding. While intrinsic NP curvature could in theory promote tube folding, we found that intrinsic curvature at the NP midline and NPBs is not sufficient for tube closure in a model adapted to measurements from our mouse live-imaging data (Fig. 3F). In contrast, a model based on notochord adhesion, mesoderm area expansion and NNE width expansion was sufficient for MHP formation and neural tube closure (Fig. 4D).

### Intrinsic versus extrinsic forces in MHP formation

Our results challenge the idea that IKNM is an active contributor to mammalian spinal NTF. The MHP contains an increased proportion of wedge-shaped cells with basally located nuclei, which has led to the view that IKNM is a driving force in avian and mammalian NTF [14, 16]. However, the increase in wedge-shaped cells at the MHP does not make implications about the cause of wedging. As a tube inner circumference can only be smaller than the tube outer circumference, it follows that neuroepithelial cells have to show increased wedging at strongly curved regions. This is independent of whether wedging is an active neuroepithelium-intrinsic process driving NTF, or a reactive process resulting from tissue deformation by external forces. Importantly, a human ectodermal *in vitro* model of neural tube folding does not form a MHP in the absence of mesoderm at physiologically representative neural tube widths [40], indicating that external tissues may be required for MHP formation. Our computational model indicates that neuroepithelial cell wedging could occur as a result of mesoderm and NNE expansion combined with NP anchoring to the notochord (Fig. 4D). In this scenario, MHP nuclei could be pushed into a basal localization through mesoderm-driven neuroepithelial cell wedging. Future research should clarify this through experimentally manipulating the mesoderm volume in developing mouse embryos or *in vitro* NTF model systems. We predict that mesoderm volume increase during folding will lead to increased MHP curvature and increased basal localization of neuroepithelial nuclei at the MHP.

### Biomechanical mechanism of DLHP formation

Our findings suggest that DLHPs emerge as a result of zippering in regions with low mesoderm expansion (Fig. 5). In these regions, the NP lateral-basal sides elevate above the mesoderm and adhere to the NNE. This could make the NP lateral sides more mobile than the medial NP, which is attached to the underlying mesoderm and notochord [41]. Thus, while zippering pulls the NP lateral tips towards the dorsal midline, the medial NP remains attached to the underlying tissue. As a result, the NP bends at the mesoderm/NNE boundary (Fig. 5E). In this framework, DLHP formation is a passive reaction to zippering rather than a process which actively contributes to neural tube closure. If zippering drives DLHP formation, laser ablation at the point of zippering should reverse dorsolateral curvature, which indeed appears to have been shown in a previous study [7].

### Molecular mechanism of DLHP formation

Our mouse and human embryo data show that mesoderm area decreases between early and late-stage spinal NTF whereas neural plate width increases (Fig. 3A,B, Fig. 4A,B, Fig. 5B,C). This shift in the neural-to-mesodermal tissue balance drives the transition from late Mode 1 (MHP) to Mode 3 (DLHPs) folding in our computational model. During posterior body formation, the neural-to-mesodermal cell ratio is regulated by bipotent neuromesodermal progenitor cells (NMPs) which can differentiate into both paraxial (presomitic) mesoderm and neuroepithelial cells [42, 43]. Termination of body axis elongation is characterized by NMP depletion and high NMP differentiation into neural tissue at the expense of paraxial mesoderm [44, 45]. NMP proliferation and differentiation depend on several molecular pathways, in particular WNT, FGF, and retinoic acid (RA) signaling. Interestingly, these pathways have also been implicated in neural tube folding and hingepoint formation. WNT and FGF signaling together drive NMP self-renewal as well as NMP differentiation towards paraxial mesoderm [46–48]. *Wnt3a*^*-/-*^ knockout mice show fewer mesoderm cells and more neural cells compared to wildtype embryos, and appear to show early DLHPs at E9 and an early transition to Mode 3 folding at E9.5 [48–50]. A similar phenotype appears to be present in *β-catenin* conditional knockouts as well as *Fgf4* /*Fgf8* conditional double knockout mice [51, 52]. RA counteracts WNT and FGF signaling by promoting NMP differentiation into neural cells [53]. *Raldh2*^*-/-*^ mouse embryos, which lack RA synthesis, show decreased neural cell identity and increased mesoderm tissue in the caudal lateral epiblast and appear to show a stronger MHP during early NT folding between E8–E8.5 [54, 55]. Inversely, embryos treated with RA during NT folding appear to have less mesoderm tissue and show sharper DLHPs [56].

A previous study found that BMP signaling blocks DLHP formation, with *Bmp2*^*-/-*^ mutants showing premature appearance of DLHPs at SS9 [11]. The mechanism through which BMP-2 exerts this effect remains unknown. Loss of BMP signaling causes ectopic neural induction and loss of mesodermal cell identity [57]. Additionally, dual-SMAD inhibition promotes differentiation of human ES, iPS and NMP cells into neural cells *in vitro* [58, 59]. In line with our model, it would be interesting to study if BMP signaling blocks DLHP formation through shifting the neural-to-mesodermal cell ratio towards more mesoderm.

To conclude, our results suggest hingepoint formation is not a driving force in spinal NTF but can rather be viewed as a folding side effect. Transition from Mode 1 to Mode 3 folding is directly linked to a shift from mesodermal to neural tissue in our model. NTF mutants with aberrant hingepoints may therefore best be understood in the context of PNP neural/mesodermal tissue dimensions.

## 4 Methods

### Computational model

The folding neural tube was modeled as an expanding transverse ectoderm cross section in 2D. We minimized the two-dimensional elastic energy density

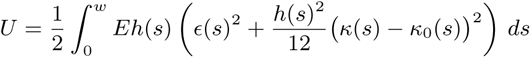

with the finite element method. Energy density refers to energy per unit length, where the length dimension is the rostrocaudal axis perpendicular to the simulated cross section. The ectoderm thickness *h*(*s*) is a field that depends on the location given by the arclength parameter *s* as detailed below. Young’s modulus *E* sets the energy density scale, equally affecting all deformation modes. *ϵ* is the tensile Cauchy strain, *κ* the curvature along the tissue centerline, and *κ*_0_ = *κ*_int_ + *κ*_p_ the curvature contribution from intrinsic curvature and plastic stress relaxation. We employed third-order beam elements containing an additional energy term for transverse shear that depends on Poisson’s ratio *ν*. A detailed description can be found in [32]. A predictor-corrector method from the Newmark family was used to integrate Newton’s equations of motion in time, using a uniform homogeneous mass density *ρ* in the tissue. For numerical equilibration, we used a damping density *γρ*, where *γ* defined the viscous relaxation rate of the tissue.

Simulation parameters are listed in Table 1. The tissue was assumed to have homogeneous and uniform material properties both within and between NP and NNE. Simulations started from a horizontal configuration unless otherwise specified. A small random spatial perturbation was imposed on the ectoderm to allow for symmetry breaking through Euler buckling and to allow for independent simulation repetitions. Between 150– 500 elements were used to discretize the ectoderm, with higher element densities applied at thin tissue regions and regions with a high width expansion. At the ectoderm end points, displacement was fixed but rotations were unconstrained (pinned boundary conditions). Simulations were rendered using ParaView 5.5.0.

**Table 1:** Model parameters

| Symbol | Value | Dimension | Description |
| --- | --- | --- | --- |
| $E$ | 1 kPa | mass/length×time<sup>2</sup> | Young’s modulus of the tissue |
| $\nu$ | 0.5 | — | Poisson’s ratio of the tissue |
| $\rho$ | 1 g/cm<sup>3</sup> | mass/length<sup>3</sup> | Mass density of the tissue |
| $K$ | 10$E$ | mass/length×time<sup>2</sup> | Bulk modulus of the mesoderm |
| $H$ | 0.05$E$ | mass/length×time<sup>2</sup> | Kinematic hardening modulus of the tissue |
| $\sigma_Y$ | 0.005$E$ | mass/length×time<sup>2</sup> | Yield stress of the tissue |
| $\gamma$ | 1000 s<sup>-1</sup> | 1/time | Viscous relaxation rate of the tissue |
| $l_a$ | 3 μm | length | Spatial range of adhesion (Model II) |
| $k$ | 3 mN/m | mass/time<sup>2</sup> | Spring constant of adhesion (Model II) |
| $t$ | variable | time | Simulated time (value between 0 and 1s) |
| $h_{NP}$ | 45 μm | length | Neural plate height (apicobasal) |
| $h_{NNE}$ | 8 μm | length | Non-neural ectoderm height (apicobasal) |
| $h_{HP}$ | variable | length | Neural plate (median) hinge point height (apicobasal) |
| $w_{NP}$ | variable | length | Neural plate width |
| $w_{NNE}$ | variable | length | Non-neural ectoderm width |
| $w_{HP}$ | variable | length | Hinge point width |
| $\kappa_{int}$ | variable | 1/length | Intrinsic curvature of the ectoderm |
| $\kappa_{mid}$ | variable | 1/length | Intrinsic curvature of the ectoderm at the middle of the hinge point domain |
| $\kappa_{max}$ | 2/$h_{NP}$ | 1/length | Theoretical maximum curvature of the ectoderm |
| $d$ | variable | length | Distance between the middle of the hinge point domain and the NP midline |
| $\alpha$ | variable | — | Mesoderm area increase factor |

### Tissue growth

Volumetric growth of the tissue was modeled by expanding the reference length of each finite element over time according to a growth multiplier *g*. For Model I, *g*(*t*) = 1 + *t*. See [60] for implementation details.

### Stress relaxation

To model stress relaxation, we used a bi-linear elasto-plastic stress-strain relationship for the bending moment in all simulations, with linear kinematic hardening and a rate-independent associative flow rule, as developed in [60]. The resulting bending moment density reads

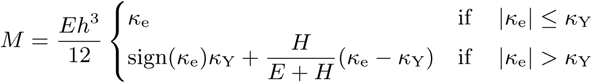

where *κ*_e_ = *κ*−*κ*_0_ denotes the elastic curvature, and *κ*_Y_ = 3*σ*_Y_*/Eh* is the effective curvature at the yield point, defined by the yield stress *σ*_Y_. In order to let almost all bending stress dissipate, both the yield stress *σ*_Y_ and the kinematic hardening modulus *H* were kept small (Table 1).

### Midline thinning

The height transition from the lateral NP to the NP midline was modeled using a smooth function:

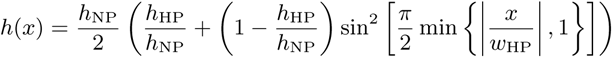

where *h*_NP_ is the lateral NP height, *h*_HP_ is the (median) hinge-point height, and *w*_HP_ is the hingepoint domain width.

### Intrinsic curvature

Intrinsically curved domains were modeled using a smooth function with the highest intrinsic curvature in the domain center:

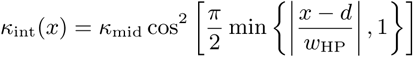

with *κ*_mid_ the intrinsic curvature at the HP domain center. *d* denotes the distance from the HP domain center to the NP midline.

### Adhesion and contact

In Model II, adhesion between adjacent NP and NNE was modeled with a bi-linear traction-separation law. Given a separation *l* between the two tissue surfaces, an adhesive force

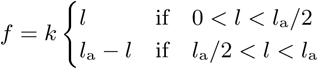

was applied, where *k* is the adhesive spring constant. In the case of volumetric overlap (*l <* 0), a linear repulsive Hertzian contact force was applied.

### Neural plate borders

For all simulations including both NP and NNE, NPB zones were modeled for the domain *w*_NP_*/*2−*w*_NPB_ *<*|*x*|*< w*_NP_*/*2 using a smooth tissue height transition:

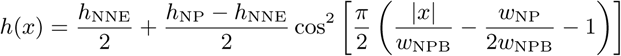

with neural plate border zone width *w*_NPB_ = 25 µm.

### Model I

In Model I, an integral constraint preserved the area underneath the ectoderm during ectoderm width expansion, using a bulk modulus *K* as Lagrange multiplier. Model I simulations used a neural plate height *h*_NP_ = 45 µm and an initial epithelium width *w*_NP_ = 500 µm. For simulations including NNE, the NP width was set to *w*_NP_ = 250 µm. NNE width was set to *w*_NP_ = 125 µm on both sides and the NNE height was fixed at *h*_NNE_ = 8 µm.

### Model II

In Model II, the target value in the area constraint increased linearly with time to simulate mesoderm expansion. The mesoderm expansion to the left and the right of the tissue midline was constrained independently. The ectoderm midline was fixed at its initial coordinates to mimic NP-notochord adhesion. The NP height was set to *h*_NP_ = 45 µm and the NNE height to *h*_NNE_ = 8 µm. NP and NNE adhered to each other when reaching a proximity of *l*_a_ = 3 µm. To promote NP-NNE adhesion, intrinsic curvature was incorporated at the NPBs (*κ*_mid_ =−*κ*_max_, *w* = 3*h*_NNE_). Model II included midline tapering with *h*_NP_*/h*_HP_ = 2 and *w*_HP_ = 2*h*_NP_, unless specified otherwise (Fig. 4G).

### Closure 1 simulations

Closure 1 (Fig. 4) was simulated using Model II. The simulation started in a concave configuration defined by 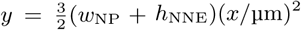. Closure 1 simulations had an initial epithelium width *w*_NP_ = 250 µm. The mesoderm area increased linearly over time from *A*_meso_(*t* = 0) = 0 µm^2^ to *A*_meso_(*t* = 1) = 1.7*×*10^4^ µm^2^. The NNE started at width *w*_NNE_(*t* = 0) = 20 µm on both sides and linearly increased over time to *w*_NNE_(*t* = 1) = 100 µm.

### PNP simulations

NTF in the PNP was simulated using Model II, starting out from a horizontal configuration with 100 µm NNE on both sides of the NP. For the simulations in Fig. 5A, the NP width was set to *w*_NP_ = 400 µm. Zippering was included by letting the NPBs converge linearly towards the midline along the *x*-axis (while keeping NPB movement along the *y*-axis unconstrained), reaching *x* = 0 at *t* = 1. Mesoderm area increased linearly over time, proportional to a half ellipse:

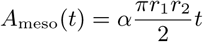

with *r*_1_ = *w*_NP_*/*2 + *w*_NNE_(0) − *h*_NP_*/*2, *r*_2_ = *w*_NP_*/*2 and *α* the mesoderm area expansion factor. NNE width was increased linearly over time and uniformly in space on each side of the PNP to match the corresponding circumference of the half ellipse:

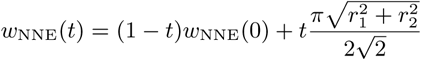

Note that *w*_NNE_(*t*) increases with *w*_NP_, but is independent of *α*.

### Microscopy data analysis

Image measurements were performed in ImageJ 1.52r-1. Neural plate apicobasal height was measured at the NP lateral side midpoint. NNE apicobasal height was measured between 5–30 µm of the NP-NNE domain border. NNE width was measured at one of the embryo lateral sides as either the NNE length from the NP-NNE domain border to the border between NNE and extra-embryonic tissue (mouse live imaging data) or as the NNE length along the embryo area dorsolateral to the notochord (in the mouse PNP and human data). Mesoderm area was measured as either the total mesoderm area at one of the embryo lateral sides (mouse live imaging data) or by measuring the mesoderm area dorsolateral to the notochord at one of the embryo lateral sides (in the mouse PNP and human data).

### Mouse live images

CAG-tdTomato-2A-H2B-EGFP (CAGTAG1) mice expressing ubiquitous membrane and nuclear markers were acquired from [61]. Lifeact-mRFPruby mice (which visualize F-actin [62]) were obtained from [63]. All mice were maintained on a mixed, CD-1/129Sv/C57BL/6J background. Developing E6.5–E8.5 mouse embryos were imaged using a light-sheet microscope for adaptive imaging of mouse embryo development. Embryos were prepared, cultured and imaged as described in [33]. Parameters were measured at every somite and at 3 time points during folding progression: “Start”, “Elevated”, and “Closure”. These time points were different for each somite. “Start” refers to the time point after gastrulation where the NP has become distinguishable from the NNE. “Closure” is the first time point where the lateral sides of the neural plate touch at the dorsal midline. “Elevated” refers to the middle time point between “Start” and “Closure”.

### Mouse PNP section data

Mouse embryo images analyzed in this study were obtained from the EMAP eMouse Atlas Project [38] and Deciphering the Mechanisms of Developmental Disorders (DMDD) [8]. DMDD is a program funded by the Welcome Trust with support from the Francis Crick Institute. The original data and image specifications can be found on the EMAP and DMDD websites, and usage of both data sources is licensed under a Creative Commons Attribution license. From these online databases, all wildtype embryos in the stage of PNP folding were included (data accessed November 2020). An overview of mouse embryos used is given in Table S2. In total, 5 Mode 1 embryos (SS11– SS22), 13 Mode 2 embryos (SS19–SS27) and 7 Mode 3 embryos (SS25–SS31) were analyzed. HREM data (DMDD) and high-resolution plastic section data (EMAP) were processed in Imaris 9.5.1 using an oblique slicer to obtain images perpendicular to the posterior neuropore. Resulting images were further analyzed in ImageJ 1.52r-1.

### Human embryo section data

Human embryos included in this study are historical specimens from the Carnegie collection (Silver Spring, MD, USA) and the Boyd collection (University of Cambridge, UK). Transverse section images of historical embryo specimens were obtained from the Virtual Human Embryo Project [34] and the 3D Atlas of Human Embryology [35]. Specimens included for analysis were between SS4 (the earliest stage embryo available which showed a folding NT) and SS13 (the earliest stage embryo available which showed complete NT closure between somite positions 1–8). Specimens 3709 (SS4), 6330 (SS7) and 6344 (SS13) originated from the Carnegie collection. Specimen H712 (SS6) originated from the Boyd collection. An overview of specimen origins and section specifications is published in [35].

## Acknowledgements

This work was funded by Swiss National Science Foundation Sinergia grant CRSII5_170930, with additional funding by a Utrecht Selective Life Sciences ExtraCurricular Track travel grant (Utrecht University, The Netherlands) and a Swiss-European Mobility Programme scholarship. The computational model was developed with partial funding by ETH Zürich under ETH Independent Investigators’ Research Awards grant no. ETH-03 10-3. We thank Laura Schaumann for technical assistance, and James Briscoe and Lisa Conrad for valuable discussions.

## Author Contributions

RV & DI conceived the study. VdG developed the different NP folding models, performed the computer simulations, analyzed the images, and produced the figures. VdG & RV produced the simulation movies. RV developed the computational framework. KMD processed the mouse live imaging data and produced the mouse live-imaging movie. All authors contributed to the writing of the manuscript.

## Supplementary Information

**Movie S1**. Model I simulation with basic model conditions.

**Movie S2**. Model I simulation with midline tapering (*h*_NP_*/h*_HP_ = 2 and *w*_HP_ = 2*h*_NP_).

**Movie S3**. Model I simulation with midline intrinsic curvature (*κ*_mid_ = *κ*_max_ and *w*_HP_ = 2*h*_NP_).

**Movie S4**. Model I simulation with dorsolateral intrinsic curvature (*κ*_mid_ = *κ*_max_*/*2, *w*_HP_ = 2*h*_NP_ and *d* = 2*h*_NP_).

**Movie S5**. Model I simulation combining midline tapering and midline intrinsic curvature (*h*_NP_*/h*_HP_ = 2, *κ*_mid_ = *κ*_max_ and *w*_HP_ = 2*h*_NP_).

**Movie S6**. Model I simulation as in Movie S5 with a NNE thickness profile (*h*_NNE_ = 8 µm, *w*_NNE_ = 125 µm on both sides, *w*_NP_ = 250 µm).

**Movie S7**. Model I simulation as in Movie S6 with differential width expansion. NNE expands in width while NP width stays constant. Final total tissue width is 1000 µm.

**Movie S8**. Model I simulation as in Movie S7 with NPB intrinsic outward curvature (*κ*_mid_ = −*κ*_max_ and *w*_BHP_ = 2*h*_NP_).

**Movie S9**. Mouse live imaging of early neural tube closure at E8.5, somite position 4.5, transverse view. Magenta, cell membrane; white, nucleus.

**Movie S10**. Model II simulation mimicking Closure 1 (concave initial configuration). Green, NNE; Blue, NP.

**Movie S11**. Model II simulation mimicking Closure 1 (concave initial configuration), without midline tapering. Green, NNE; Blue, NP.

**Movie S12**. Model II simulation mimicking folding in the posterior neuropore (flat initial configuration), without zippering, mesoderm area increase factor *α* = 1. Green, NNE; Blue, NP.

**Movie S13**. Model II simulation mimicking folding in the posterior neuropore (flat initial configuration), without zippering, mesoderm area increase factor *α* = 2. Green, NNE; Blue, NP.

**Movie S14**. Model II simulation mimicking folding in the posterior neuropore (flat initial configuration), with zippering, mesoderm area increase factor *α* = 1 (Mode 1). Green, NNE; Blue, NP.

**Movie S15**. Model II simulation mimicking folding in the posterior neuropore (flat initial configuration), with zippering, mesoderm area increase factor *α* = 0.625 (Mode 2). Green, NNE; Blue, NP.

**Movie S16**. Model II simulation mimicking folding in the posterior neuropore (flat initial configuration), with zippering, mesoderm area increase factor *α* = 0.125 (Mode 3). Green, NNE; Blue, NP.

**Table S1:** Mouse live imaging measurements per somite position and folding stage

| Somite position | Folding stage | $N$ |
| --- | --- | --- |
| 1 | start | 2 |
| 1 | elevated | 1 |
| 1 | closure | 1 |
| 2 | start | 3 |
| 2 | elevated | 2 |
| 2 | closure | 2 |
| 3 | start | 3 |
| 3 | elevated | 3 |
| 3 | closure | 3 |
| 4 | start | 3 |
| 4 | elevated | 3 |
| 4 | closure | 3 |
| 5 | start | 3 |
| 5 | elevated | 3 |
| 5 | closure | 3 |
| 6 | start | 3 |
| 6 | elevated | 3 |
| 6 | closure | 3 |
| 7 | start | 2 |
| 7 | elevated | 2 |
| 7 | closure | 2 |
| 8 | start | 2 |
| 8 | elevated | 1 |
| 8 | closure | 1 |

**Table S2:** Overview of mouse specimens used for posterior neuropore analysis

| Origin | embryo ID | Somite stage (estimate) | Folding mode |
| --- | --- | --- | --- |
| EMAP | EMA:13 | 11 | 1 |
| EMAP | EMA:14 | 17 | 1 |
| DMDD | DMDD1835 | 20 | 1 |
| EMAP | EMA:36 | 22 | 1 |
| EMAP | EMA:37 | 22 | 1 |
| DMDD | DMDD1834 | 19 | 2 |
| DMDD | DMDD3544 | 20 | 2 |
| EMAP | EMA:28 | 22 | 2 |
| DMDD | DMDD3527 | 24 | 2 |
| DMDD | DMDD3541 | 24 | 2 |
| DMDD | DMDD3528 | 25 | 2 |
| DMDD | DMDD3532 | 25 | 2 |
| DMDD | DMDD3500 | 26 | 2 |
| DMDD | DMDD3510 | 27 | 2 |
| DMDD | DMDD3514 | 27 | 2 |
| DMDD | DMDD3521 | 27 | 2 |
| DMDD | DMDD3524 | 27 | 2 |
| DMDD | DMDD3538 | 27 | 2 |
| DMDD | DMDD3517 | 25 | 3 |
| DMDD | DMDD3507 | 26 | 3 |
| DMDD | DMDD3536 | 26 | 3 |
| DMDD | DMDD3531 | 27 | 3 |
| DMDD | DMDD3506 | 28 | 3 |
| DMDD | DMDD1904 | 31 | 3 |
| DMDD | DMDD1911 | 31 | 3 |

